# Kinetic relaxation of giant vesicles validates diffusional softening in a binary lipid mixture

**DOI:** 10.1101/2022.09.28.509943

**Authors:** Kayla Sapp, Mina Aleksanyan, Kaitlyn Kerr, Rumiana Dimova, Alexander Sodt

## Abstract

The stiffness of biological membranes determines the work required by cellular machinery to form and dismantle vesicles and other lipidic shapes. Model membrane stiffness can be determined from the equilibrium distribution of giant unilamellar vesicle surface undulations observable by phase contrast microscopy. With two or more components, lateral fluctuations of composition will couple to surface undulations depending on the curvature sensitivity of the constituent lipids. The result is a broader distribution of undulations whose complete relaxation is partially determined by lipid diffusion. In this work, kinetic anaysis of the undulations of giant unilamellar vesicles made of phospatidylcholine-phosphatidylethanolamine mixtures validates the molecular mechanism by which the membrane is made 25% softer than a single-component one. The mechanism is relevant to biological membranes, which have diverse and curvature-sensitive lipids.

## I. INTRODUCTION

The cellular membrane is a complex mixture of many lipids and proteins, which may be attached peripherally, reside in one leaflet, or cross both. Membrane shape is highly influenced by a complicated cytoskeletal network. To isolate the mechanical effect of individual lipid components in such a system is currently infeasible. Giant unilamellar vesicles (GUVs) are an excellent membrane model system to which complexity can be introduced gradually [1]. Rather than attempting to visualize the distributions and motions of individual membrane components spectroscopically, with GUVs the influence of those components on the projected bilayer shape can be directly observed.

GUV mechanics are typically described using the Helfrich/Canham [2, 3] (HC) energy density, *H*:

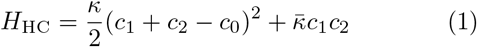

where *κ* is the membrane bending rigidity, *c*_1_ + *c*_2_ is the total curvature where *H*_HC_ is being evaluated, *c*_1_*c*_2_ is the Gaussian curvature, and *c*_0_ is the bilayer spontaneous curvature.

We refer to the range of the magnitude of undulations of the GUV as the dynamic ensemble and to the relaxation times of the undulations as GUV kinetics. GUV mechanics are typically analyzed in terms of the dynamic ensemble. The bending stiffness is typically inferred from the range of the dynamic ensemble; larger fluctuations indicate a softer bilayer susceptible to thermal agitation. The stiffness also impacts kinetics; with a stronger restoring force, stiffer bilayers relax more quickly. The undulation of a GUV is visible to both phase contrast and confocal microscopy [4].

Some membrane mechanical parameters can also be inferred from static structures under external stress. An estimate of the stiffness of the red blood cell [5] as well as simple model membranes [6] can be obtained from analysis of the shape of membranes under micropipette aspiration. Even the challenging modulus of Gaussian curvature can be deduced from the analysis of the shapes of GUVs composed from a ternary mixture that phase separate into microscopic ordered and disordered domains [7]. In this case, the external stress is the line tension between domains, to which the shape of the surrounding vesicle adapts. The spontaneous curvature of lipid constituents can also be inferred by pulling nanoscale tubes using optically-trapped beads. The force required for inward or outward tubulation will depend on the spontaneous curvature of the whole bilayer [8–10].

Setting aside the extreme case of macroscopic phase separation, lipid *mixtures* of complex molecular composition raise the possibility of lipid-lipid interactions giving rise to inhomogeneity invisible to microscopy. When nanometer-scale heterogeneity is a strong determinant of mechanical properties, the variation of lipid concentrations will modify the fluctuation spectrum of GUVs non-linearly. The characterization of the effect of complex lipid heterogeneity by GUV fluctuations is closely related to the main target of interest: a model of the mechanics of cellular membranes.

This work examines a simple mechanism of softening in complex membranes, what we term *diffusional* softening [11]. Diffusional softening results from the dynamic coupling between the lateral distribution of lipids and the membrane undulations. It has been proposed as a method for determining lipid or protein diffusion constants [12–14]. The mechanism only applies for leaflets with a mixture of lipids with varied spontaneous curvature. It is independent of the asymmetry of composition between leaflets. The effect was originally described by Leibler in 1986 for general inclusions [15], likely applicable to inclusions like alamethicin [16], fusion peptides [17] and other proteins [14], but the theory applies equally well to lipids [12, 18–20]. The width and relaxation time of nanotubes of lipid mixtures pulled from black lipid membranes strongly imply that lipid sorting leads to constriction of the tube, implying softness [21]. Non-linear variation of *κ* has been observed in simple simulations of two component mixtures, in which the mixture of a stiff and soft lipid appears softer than even a pure bilayer of the soft lipid (see Fig. 3a of Ref. [22] and Fig. 8 of Ref. [23]). As shown below, the softening of *κ* is quadratic in the spontaneous curvature difference between lipids, and goes as χ(1–χ), where χ is the mixed mole fraction for a binary mixture, consistent with the observations in Refs. [22, 23].

Whereas the undulations of a single component bilayer relax with timescale 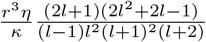, for diffusional softening the timescale is perturbed by the relaxation of the lateral compositional fluctuation, which goes as 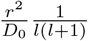. Here *η* is the solvent viscosity, *l* is the degree of the spherical harmonic (SH), *r* is the vesicle radius, and *D*_0_ is the diffusion constant. This mechanism can thus be distinguished by the *l*-dependence of the undulation relaxation time. The time-scales also differ in their dependence on the vesicle radius.

The bending modulus for gel-phase systems approaches zero near the phase transition (see Fig. 13 of Ref. [24] and Fig. 6 of Ref. [25]). Thermal fluctuations of many mechanical properties are sufficient to explain this, including the area-per-lipid, which differs greatly between the gel and fluid phases. Differing curvature preference of gel and fluid phases would also contribute to bilayer softening. This is a clear indication of how dynamically fluctuating material properties lead to profound changes in softness. Moreover, the kinetics of the gel-fluid transition should influence the relaxation time of visible undulations.

This work uses analysis of the kinetics of GUV relaxation, supported by simulation, to distinguish the intrinsic bending modulus from the diffusionally-softened bending modulus.

The dynamic fluctuations of GUVs composed either completely of POPC or of 40% DOPE and 60% POPC are first presented. The reduced softness of the mixed system (22*k*_B_*T*) compared to pure POPC (28*k*_B_*T*) is shown to be consistent with diffusional softening on the basis of a kinetic fit to the autocorrelation function of the GUV undulation amplitude.

To validate the model, the kinetics of the relaxing GUV are compared to continuum simulations that incorporate the HC energy, as well as the experimentally-validated relaxation times of the undulation and lateral-compositional fluctuations. Fitting the time-dependence of auto-correlation functions is not straightforward. Choices of how to compare fits to the experimental data impact the accuracy and precision of the extracted model parameters. To account for this ambiguity in an even-handed way, the expected error and optimal fitting strategy is derived from the simulations, rather than the experiments. Then, the fitting scheme is applied to the time auto-correlation function of GUV undulations, as measured by phase-contrast microscopy.

## II. METHODS

The theory is first developed to describe the mechanism of diffusional softening, including the kinetics necessary for modeling. The experimental and simulation protocols for characterizing GUV fluctuations are then provided, as well as how a framework was developed to most precisely fit the experiment as well as to anticipate stochastic error.

### A. Theory

The HC energy density is modified by the presence of PE by subtracting the homogeneous (background) HC energy and adding in the contribution from the PE lipid:

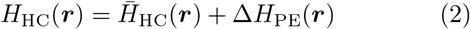

where

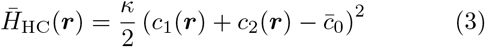

and

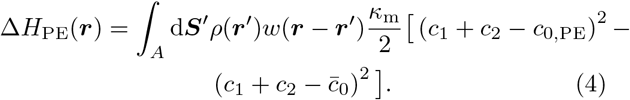

Here *w*(***r***) is the spatial extent of a single lipid [26] and *ρ*(***r***′) is the number of PE lipids per unit area in the outer leaflet of the GUV (a trivial extension to both leaflets is made below). Note that the bending modulus is assumed to be homogeneous; the change in energy density only reflects changes in *c*_0_. Assuming perfectly local spatial extent for a lipid,

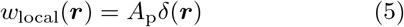

where *A*_p_ is the area of a PE lipid, yields

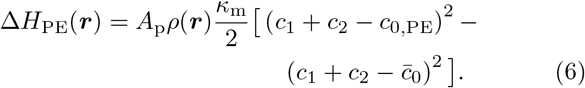

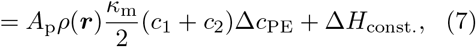

where here Δ*H*_const_. is a constant term independent of vesicle curvature. The assumption of perfectly local extent is justified as long as the undulation wavelength considered is much greater than the mechanical extent of the lipid. According to the mechanical extent of simulated PE lipids [26], this is easily justified. Larger lipidic patches requiring treatment at higher *q* could be described with finite spatial extent.

The average elastic curvature energy, without coupling to PE, is then:

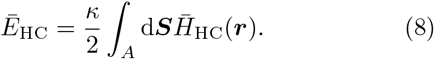

For a vesicle, both the membrane shape *R*(*θ, ϕ*) and lipid distribution *ρ*(*θ, ϕ*) are expanded in SH:

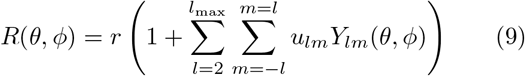

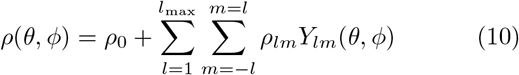

with coefficients *u_lm_* (unitless) and *ρ_lm_* (units per area). With the curvature written as the divergence of the normal, the compositionally-averaged elastic energy 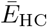 is evaluated from 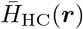 (Eq. 3) as:

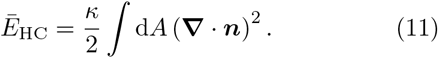

Taking a second order approximation the curvature energy (Eq. 8) is:

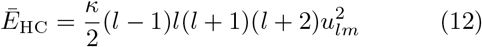

Given a bilayer with a mole fraction χ of one lipid (here DOPE) and 1 – χ background lipids (here, POPC), the energy of a density fluctuation is given by

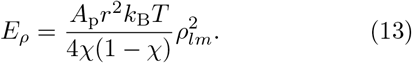

This is a purely entropic factor.

The coupling of *ρ_lm_* and *u_lm_* by Δ*H*_PE_(***r***) is similarly evaluated in SH:

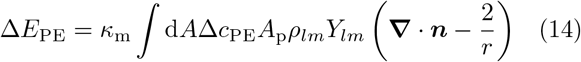

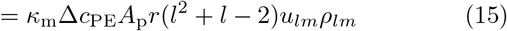

Note that in our model, Δ*E*_PE_ for the inner leaflet of the bilayer requires only switching the sign of curvature; it is the negative of Δ*E*_PE_ for the outer leaflet. As the coupling depends linearly on *ρ_lm_*, it is irrelevant whether lipids are modeled to be in the outer leaflet (with Δ*c*_PE_) or in the inner leaflet (with – Δ*c*_PE_). We therefore state, without loss of generality, that they are distributed symmetrically throughout the bilayer as in the experiment. To further cast the model as the bilayer coupling, we reintroduce the bilayer *κ*, with *κ* = 2*κ*_m_:

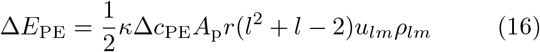

Combining Eqs. 12, 13, and 16, the total energy is

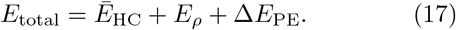

The expectation of 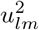 is determined by

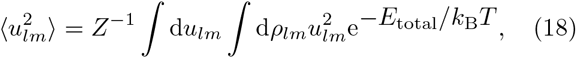

where

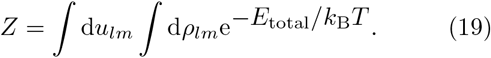

For a single-component membrane (χ = 0 or Δ*c*_PE_ = 0), integration leads to

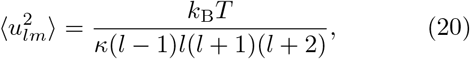

from which the bending rigidity can be determined.

At higher *q* where diffusion is slower than the relaxation of undulations, the membrane auto-correlation function reflects the time-scales of the two processes and how they are coupled through spontaneous curvature. We can derive a theoretical prediction of the autocorrelation function from the dynamics of the system.

The expectation of 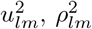, and *u_lm_ρ_lm_* are:

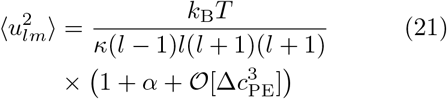

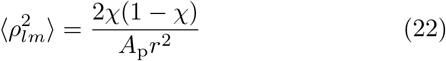

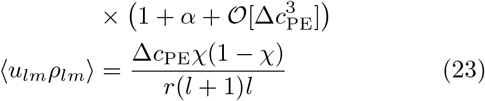

Where the softening constant

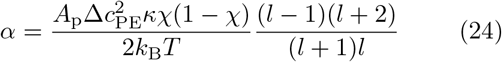

is defined for convenience, as it arises frequently.

A bilayer with a symmetric or asymmetric mixture of lipids (with unequal spontaneous curvatures) will experience apparent softening according to

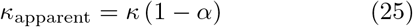

Note that the fraction depending on *l* goes to one rapidly, giving the diffusional softening for a planar system [11, 20]. This assumes that both leaflets contain the mixture χ = *ρA_p_*.

Langevin equations model the kinetic relaxation of coupled membrane undulations and lipid redistribution. They are:

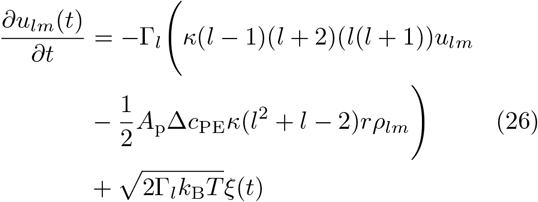

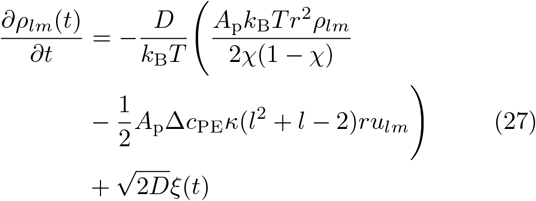

where ξ(*t*) describes a stochastic process:

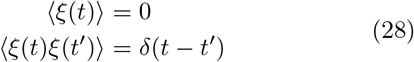

Note that when *u_lm_*(*t*) and *ρ_lm_*(*t*) are determined numerically below, ξ(*t*) is simulated by drawing random numbers from a normal distribution with zero mean and variance Δ*t*, where Δ*t* is the time-step. In Eq. 26, Γ_*l*_ describes the hydrodynamics associated with the membrane interacting with the solvent, and therefore depends on the solvent viscosity.

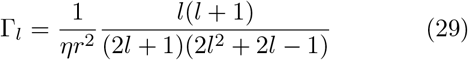

In Eq. 27, *D:*

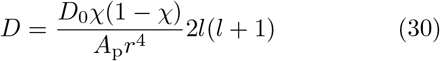

is chosen to give the appropriate diffusion time-scale. The symbolically-simplified system of two over-damped harmonic oscillators, coupled together:

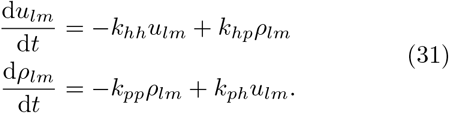

Here *k_hh_* and *k_hp_* is proportional to *η*, while *k_ph_* and *k*_pp_ are proportional to *D*. The sign of *k_hp_* is positive indicating that amplitude in *ρ_lm_* amplifies *u_lm_*; this choice implies that curvature is measured in the sense of the upper leaflet; positive *ρ_lm_* and positive *c*_0_ imply that *u_lm_* increases. This relationship is reversed in the lower leaflet (positive *ρ_lm_* implies the lipid density is decreased in the lower leaflet).

Were curvature and diffusion independent, the two processes would have relaxation timescales:

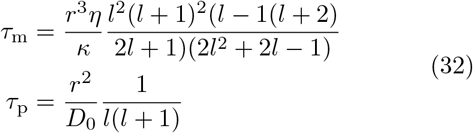

When diffusion is slower than shape fluctuations *k_pp_* << *k_hh_* and the impact of coupling to the density depends on *k_hp_* × *k_pp_* as the density slowly impacts *u_lm_*. The time autocorrelation function is:

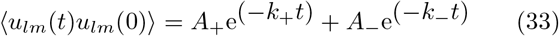

with decay constants

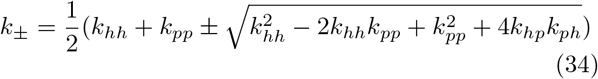

and time-scales

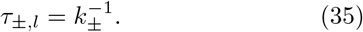

For the purpose of interpreting the different time-scales, consider *τ*_+_ as the fast membrane relaxation time-scale and *τ*_−_ as the slower diffusion time-scale. *A*_±_ are the amplitudes of each exponential.

With *k_pp_* << *k_hh_* and *k_ph_* small, to first order in the diffusion constant these rates are:

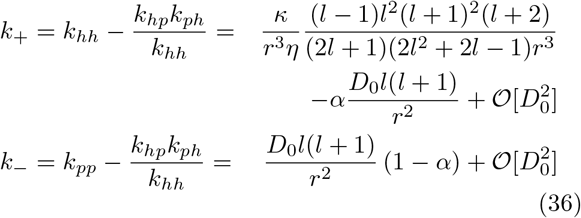

The rates are both decreased by the coupling. The amplitudes associated with each exponential decay are

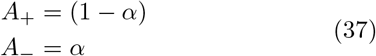

where the autocorrelation function is normalized to be 1 at *t* = 0.

### B. GUV microscopy

GUVs were prepared by the electroformation method as it is described previously [27, 28]. Briefly, pure 1-palmitoyl-2-oleoyl-sn-glycero-3-phosphatidylcholine (POPC) (Avanti Polar Lipids, Germany) or a mixture of POPC and 40 mol% 1,2-dioleoyl-sn-glycero-3-phosphoethanolamine (DOPE) (Avanti Polar Lipids, Germany) were dissolved in chloroform to a final concentration of 4 mM. Then, 10 *μ*L of the lipid solution was spread as a thin film on a pair of indium-tin oxide (ITO)-coated glass plates (PGO-GmbH, Iserlohn, Germany), which are electrically conductive. Afterwards, they were dried under a stream of Nitrogen and placed in a desiccator for 2 h to evaporate the organic solvent. A Teflon spacer with 2 mm thickness was sandwiched between the two ITO glasses (conducting sides facing each other) to form a chamber. The chamber was filled with 20 mM sucrose solution and connected to a function generator (Agilent, Waldbronn, Germany). To initiate the electroswelling process, a sinusoidal alternating current (AC) electric field at 10 Hz frequency with a 1.6 V (peak to peak) amplitude was applied for 1 hour. The obtained vesicles were harvested from the chamber and used freshly within 24 h after preparation. For fluctuation spectroscopy, the vesicle suspension was 4-fold diluted in 22 mM glucose. The osmolarity of the sugar solutions was adjusted with osmometer (Osmomat 3000, Gonotec, Germany). The vesicles were additionally deflated before imaging by leaving the observation chamber open for 5 min to let water evaporate. Membrane fluctuations were observed under a phase contrast mode of an inverted microscope, Axio Observer D1 (Zeiss, Germany), equipped with a Ph2 40 x (0.6 NA) objective. High speed video recordings were performed with a Pco.Edge camera (PCO AG, Kelheim, Germany). The image acquisition rate was set to 100 frames per second (fps) at exposure time of 200 μs. To prevent correlated images, statistics were averaged for every 4th frame. Only defect-free quasi-spherical vesicles, 8-21 *μ*m in radius and with low tension values 10^−7^ – 10^−9^ N m^−1^ were analyzed. A set of 21000 images (3× 7000 frames with 3 min gap between each recording sequence) were acquired for each vesicle. All experiments were performed at 25°C. The vesicle contour was detected through the lab owned software [29]. This included software for the construction of spatial and temporal correlation functions to characterize the shape fluctuations of the membrane. Vesicle contours were detected through the Viterbi algorithm. The amplitudes were fit with the Levenberg-Marquardt algorithm for statistical analysis and characterization of *κ*_apparent_. A χ^2^ test was applied to determine the range of modes included, with values in the range of 0.8-1.2.

### C. Simulation

Solving Eqs. 26 and 27 numerically for small discrete time increments leads to a time-series for the membrane and distribution modes. The amplitudes of undulations are projected into the equatorial plane

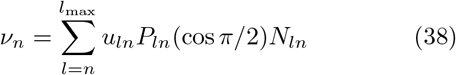

where, *P_ln_* are the associated Lengendre polynomials and 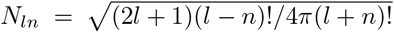 is a normalization factor. Four sets of simulations using parameters to reflect the experiments performed were done, values are in Table I. The time-step is determined by the fastest membrane undulation mode (Δ*t* << *τ*_m,min_).

**TABLE I:** Simulation parameters

|  |  |  |  |  |
| --- | --- | --- | --- | --- |
| Spontaneous Curvature, $\Delta c_{\text{PE}}$ (nm<sup>-1</sup>) | 0.0 | 0.140 | 0.280 | 0.340 |
| Surface Coverage, $\chi$ | N/A | 0.4 | | |
| Bending Rigidity, $\kappa$ ($k_{\text{B}}T$) | 33.78 | | | |
| Diffusion Constant, $D$ ($\mu\text{m}^2/\text{s}$) | N/A | 8.0 | | |
| Viscosity, $\eta$ (Pa s) | $8.9 \times 10^{-4}$ | | | |
| Area per Lipid, $A_{\text{p}}$ (nm<sup>2</sup>) | 0.634 | | | |

The undulation time autocorrelation function (〈*ν_n_*(*t*)*ν_n_*(0)〉) is calculated from the time-series of membrane undulations. This is related to the autocorrelation function of the vesicle through the projection of the average amplitudes

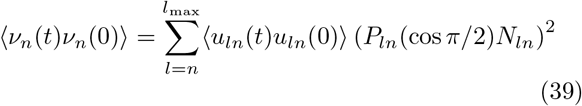

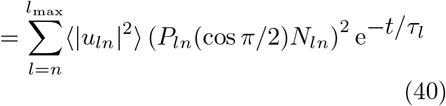

This can be fit with the analytical expression derived above in order to extract the softening factor.

An example correlation function is shown in Fig. 2, where a single short time simulation spectrum (dashed lines) is compared to the average of three simulations (solid lines). Obviously, the single simulation spectrum contains a lot of noise. The experimental spectra are also overlaid with noise. For this reason, we average over three simulations to decrease the noise in the simulation spectrum. This allows for the determination of fitting parameters that can be used with experiment.

**FIG. 1:**
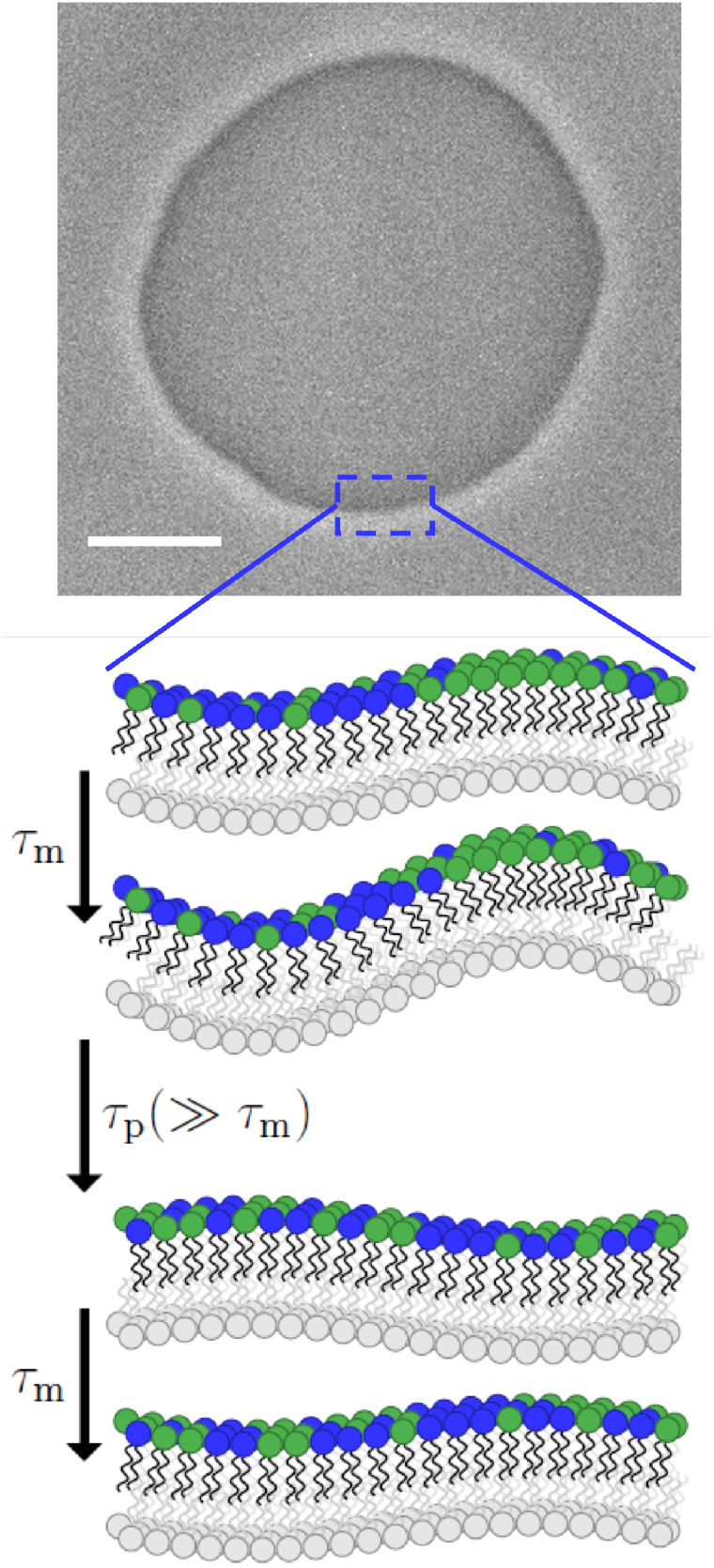
A phase contrast screenshot of a quasi spherical GUV composed of 40 mol% DOPE and 60 mol% POPC. The vesicle was prepared in 20 mM sucrose and diluted in 22 mM glucose. The scale bar is 10 μm. Zooming in a patch on the vesicle can be considered planar. Consider the green lipid to have more positive spontaneous curvature than the blue. Stochastic co-localization of the green lipid stimulates an undulation that adapts quickly (at the illustrated wavelength). Over time, diffusion relaxes the lateral distribution. The net effect is that the bilayer is softer both in appearance and practice. These fluctuations can be seen in S1A slowed-down video of a fluctuating POPC vesicle containing 40 mol% DOPE (same vesicle as in Figure 1 in the main text). The vesicle was prepared in 20 mM sucrose, 4-fold diluted in 22 mM glucose, and additionally deflated by leaving the observation chamber open for 5 minutes. The sequence shows phase contrast images acquired at 100 frames per second and displayed at 25 frames per second (processing was done with Fiji). The approximate duration is roughly 4 seconds in real time (time stamps shown in the upper left corner). **If a link to the movie is not available, please see the included material where you obtained this document.** figure.caption.3 of the Supplemental Material.

**FIG. 2:**
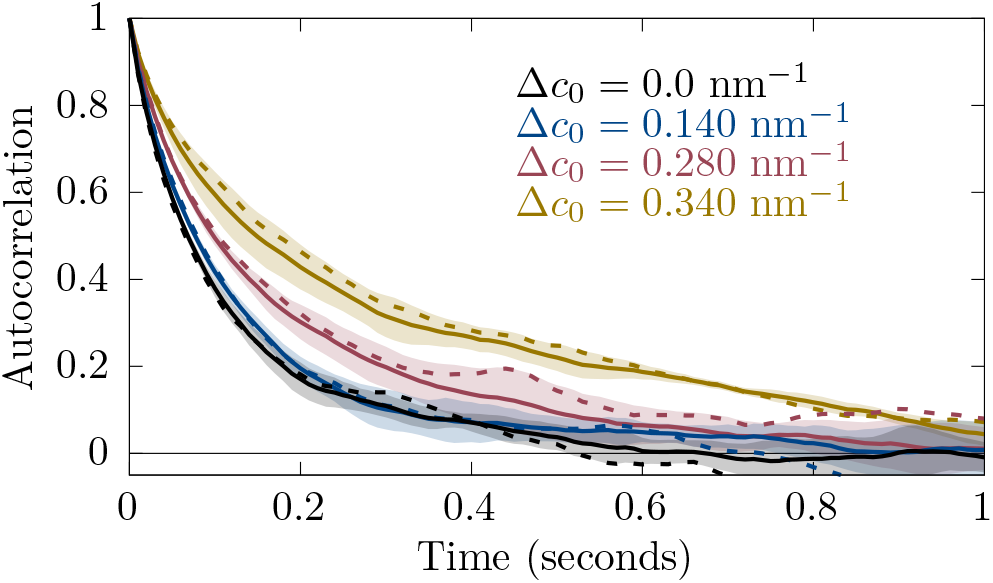
An example of autocorrelation functions from simulation (for *q* = 0.515 *μ*m^−1^). Solid lines are averaged over three runs and dashed lines are for a single run. The shaded region is the standard deviation of the average.

**FIG. 3:**
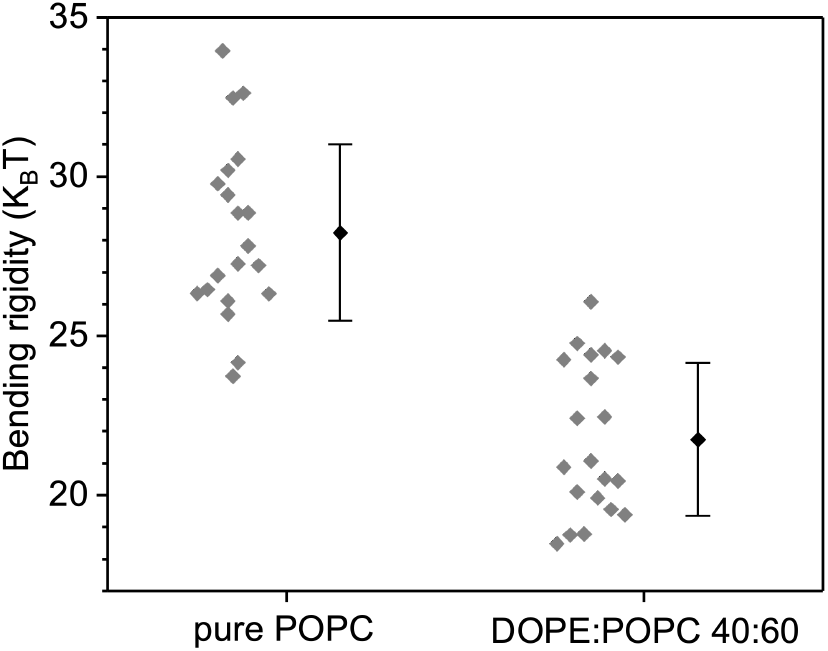
Bending rigidity of membranes made of pure POPC (100 mol%) and DOPE:POPC 40:60 mol%. Gray diamonds indicate measurements on individual GUVs. Mean and standard deviation values are shown to the right.

### D. Fitting experimental and simulation auto-correlation functions

Auto-correlation functions are fit using a functional form that replicates the theoretical dynamics implied by Eqs. 36 and 37. These relations govern the dynamics of the complete SH, yet the functional form must be for the projection into the plane, just as the simulations replicate the observable of the GUV microscopy.

Auto-correlation functions for each vesicle projection are fit by a function

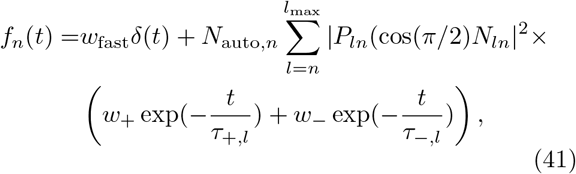

where *δ*(*t*) accounts for fast stochastic experimental noise, *τ*_+,*l*_ and *τ*_−,*l*_ are the membrane-dominated and diffusion-dominated relaxation times, respectively (Eqs. 35 and 36) while the *w* constants are weights:

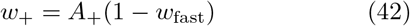

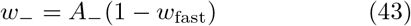

such that (with *A*_+_ + *A*_−_ = 1) the weights sum to one. The constant

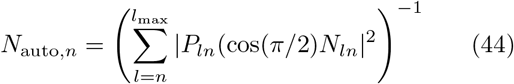

normalizes the auto-correlation function. The diffusion constant *D*, bending modulus *κ*, difference in spontaneous curvature Δ*c*_PE_ are shared parameters for the set of autocorrelation functions. Additionally, a constant modeling the magnitude of unresolvable fast (below 0.01 seconds) noise *w*_fast_ is introduced for each correlation function.

Note that *n* denotes the integer mode of the *projected* spherical harmonic, which does not decay with a single timescale even absent particle coupling. Instead, it reflects the relaxation of spherical harmonics with *l* ≥ *n*. However, the majority of the amplitude will be dominated by the lowest mode spherical harmonic with *l* = *n*. Therefore, we attach the wavenumber 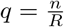 to describe autocorrelations of *ν_n_*, expecting the dynamics of modes with similar *q* but with varied *n* on vesicles of varied *R* to have similar kinetics. We use *q* to define the range of modes appropriate for fitting with the above theoretical kinetics, as well as to shift the weight bye *w_q_* of autocorrelation functions in χ^2^ to higher *q*. With their fast relaxation times, autocorrelation functions at higher *q* have compressed time domains and thus contribute weakly to χ^2^.

Optimal parameters are found by minimizing

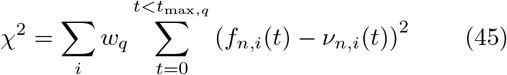

where the sum is over all autocorrelation functions for a vesicle set with *q* such that *q*_min_ < *q* < *q*_max_. Here *t*_max,*q*_ depends on *q*. The time-domain is chosen in terms of *n* multiples of the membrane relaxation time, *t*_max,*q*_ = *nτ*_m_.

We use a sum of exponentials, one with *τ*_m_ and one with *τ*_p_. The difference between these mechanisms is their *q* dependence (*q*^−3^ vs. *q*^−2^). Practically, we fit the data with the bending modulus and diffusion constants as two adjustable parameters. In the two-parameter fit each mode has an additional parameter: the weight of auto-correlation that is assigned to the diffusion mechanism.

## III. RESULTS AND DISCUSSION

The diffusional softening mechanism derived is tested on both simulation and GUV fluctuation data by fitting fluctuation autocorrelation functions to the model (Eq. 41). Fitting yields the apparent spontaneous curvature difference (Δ*c*_PE_), intrinsic bending rigidity (*κ*), and lipid diffusion constant (*D*), extracted purely from kinetics. Typically, *κ*_apparent_ (equal to (1 – *α*)*κ*) would be determined from the average fluctuations 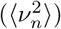. In addition to the kinetic analysis that determines *κ* and Δ*c*_PE_ and thus implies *κ*_apparent_, fluctuation analysis applied to determine *κ*_apparent_ is shown below.

### A. Apparent bending rigidity from GUV fluctuations

The classic GUV fluctuation experiment extracts the bending modulus from equilibrium fluctuations 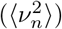:

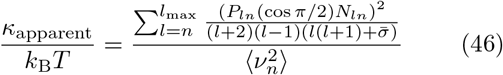

The bending modulus and tension are fit as in Ref. [29]. Following determination of each individual vesicle *κ*_apparent_ by fluctuation analysis, two sets of similar vesicles were determined, one for each composition. The range in *κ*_apparent_ for inclusion into the sets was determined by sorting the vesicles according to *κ*_apparent_ and selecting a range with minimal variation in *κ*_apparent_. A set of GUVs with similar apparent bending modulus were selected for analysis, one for PE/PC (8 vesicles with *κ*_apparent_ from 19.59 to 21.28 *k*_B_*T* with mean 20.27± 1.18 *k*_B_*T*) and one for POPC (nine vesicles with *κ*_apparent_ from 26.01 to 28.21 *k*_B_*T* with mean 26.69 ± 1.86 *k*_B_*T*). The rationale is that variations in bending modulus, for whatever reason, would imply a variation in relaxation timescale — although the timescales would still be well separated. Averaging over all the GUVs, the *κ*_apparent_ for PE/PC was 21.74 ± 2.52 *k*_B_*T* and for POPC was 28.24 ± 3.19 *k*_B_*T*, which is shown in 3.

### B. Kinetic fits to simulation

Fitting the simulation data validates the fitting software and underlying approach. Four simulations were run, with Δ*c*_PE_ = {0,0.14,0.28,0.34} nm^−1^, *κ* = 33.78 *k*_B_*T*, and *D* = 8 *μ*m^2^/s. The bending modulus is overestimated slightly versus the input parameter (33.78 – 36.49 *k*_B_*T*). The diffusion constant is 8 *μ*m^2^/s within one standard error (< 0.25 μm^2^/s). Extracted spontaneous curvatures are slightly overestimated: {0.02, 0.16, 0.30, 0.36} ±0.003 nm^−1^, in each case 0.02 nm^−1^ too high.

The autocorrelation function for each in-plane projected mode is fit to its model kinetics (Eq. 41). Averaging multiple autocorrelation functions together (for modes with similar dynamics) illustrates the separate undulation and diffusion timescales better than noisy individual fits. Average autocorrelation functions for three *q*-ranges are shown in Fig. 4, as well as the averages for the fits. The fit curves shown are solved for the Δ*c*_PE_ = 0.28nm^−1^ data set, adjusting single values of *κ*, *D*, and *c*_0_ applicable to all fit curves for a data set. For the simulation, fast noise is set to zero as this term only models experimental noise. Error bars are computed by statistical analysis of the correlation functions, assuming the same kinetics.

**FIG. 4:**
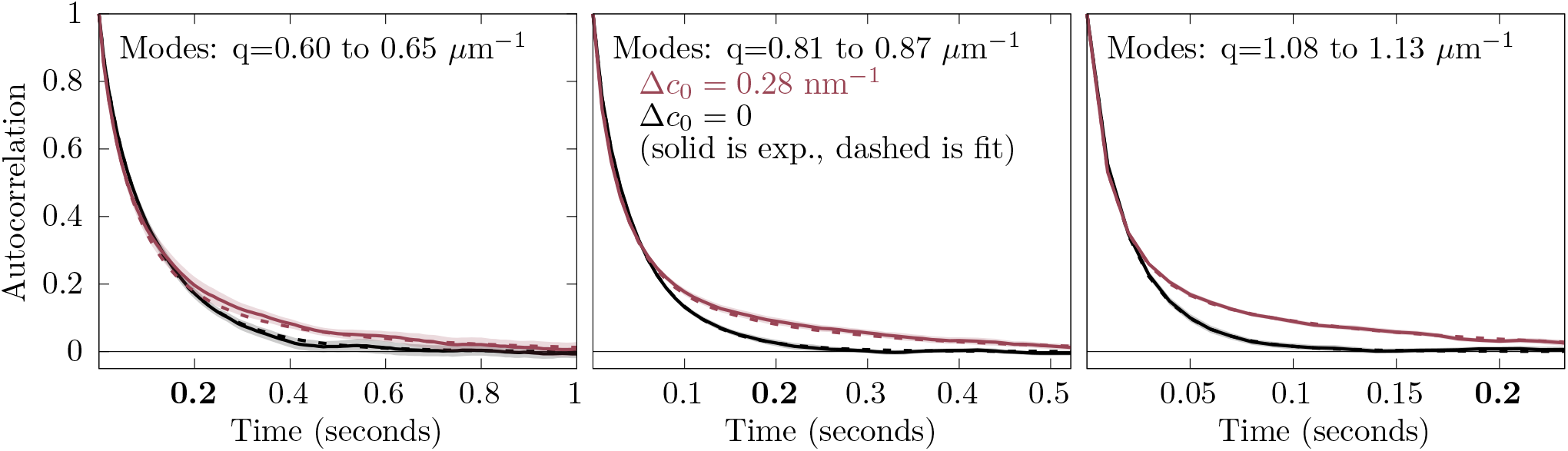
The average of the autocorrelation 〈*ν_q_*(*t*)*ν_q_*(0)〉 over similar modes for the simulation (solid) and fits (dashed). POPC is colored in black. The Δ*c*_PE_ = 0.28 nm^−1^ is colored red. The shaded regions indicate two standard errors from the simulation obtained by averaging over similar modes. The label at *t* = 0.2 s on all three plots is shown in bold for a common reference.

The difference between the red and black curves at long time illustrates the effect of curvature-coupled lipid diffusion on the undulation relaxation timescale. The agreement of the fits and input model parameters indicates that the fitting procedure is robust for simulation data with the same information content as the experiment.

### C. Kinetic fits to GUV microscopy

Average autocorrelation functions for the same three *q*-ranges above are now shown in Fig. 5 for the undulations recorded by GUV microscopy, as well as the averages for the fits. While like the simulations the single fit shares mechanical parameters between auto-correlation functions, now each auto-correlation function has its own parameterization of the the fast noise. Before plotting, the fast noise was subtracted from the autocorrelation function, the average was computed, and, finally, renormalized.

**FIG. 5:**
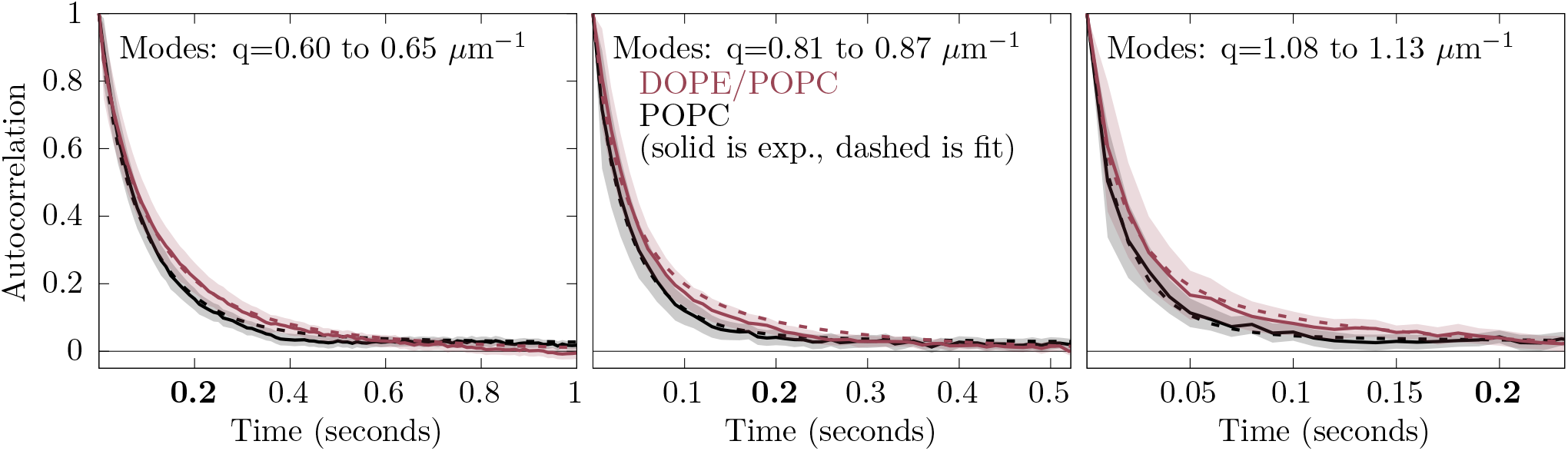
The average of the autocorrelation 〈*ν_q_*(*t*)*ν_q_*(0)〉 over similar modes for the experiment (solid) and fits (dashed). POPC is colored in black. The DOPE/POPC mixture is colored red. Filled curves indicate two standard errors from the experiment obtained by averaging over similar modes.

Fitting the mechanical parameters of the PE/PC mixture yields *κ* = 29.90 ± 1.01 *k*_B_*T*, *D* = 8.2± 0.4*μ*m^2^/s, and Δ*c*_PE_ = 0.340 ± 0.01 nm^−1^. For the fit to POPC, *κ* = 26.35 ± 0.85 *k*_B_*T*, *D* = 1.1 ± 0.2 *μ*m^2^/s, and Δ*c*_PE_ = 0.16±0.007 nm^−1^. Note that for the pure POPC GUVs, *D* and Δ*c*_PE_ should not be part of the mechanism, and therefore, ΔcPE should be zero with *D* unresolvable. We believe that, for POPC, these values are indicative of “over-fitting”, that is, the use of nonsensical parameters that fit stochastic error in the experiment and systematic error in the model. The fits indicate that, in contrast to *κ*_apparent_, *κ* is similar for the PE/PC and POPC samples. Furthermore, the difference in ΔcPE for the two samples compares well to the expected spontaneous curvature of DOPE (−0.34 nm^−1^ [30]) and POPC (−0.02 nm^−1^ [31]).

If the whole range of available GUVs were selected for analysis, the average *κ* for POPC would be 32.095 ± 1.18 *k*_B_*T* and 30.07 ± 1.52 *k*_B_*T* for PE/PC. Spontaneous curvatures were similar to the reduced set (0.18 ± 0.01 nm^−1^ and 0.30 ± 0.007 nm^−1^ for POPC and PE/PC, respectively).

The fit results converge for the experiment when *t*_max,*q*_ is sufficiently large (above 30 times *τ*_p_), and for *q*_max_ greater than 1 *μ*m^−1^. Sensitivity of *c*_0_ to *t*_max,*q*_ and *q*_max_ are shown in Figs. S3Variation in the apparent spontaneous curvature difference with *t*_max_. figure.caption.6 and S2Variation in the apparent spontaneous curvature difference with *q*_max_. figure.caption.5 of the Supplemental Material.

The fit to the pure POPC shows slow timescale relaxations; yet these are inconsistent with diffusional softening. Foremost, the apparent diffusion constant extracted (1.1*μ*m^2^/s) is inconsistent with lipid diffusion, which is consistently measured to be much larger (ca. 8*μ*m^2^/s and modestly reduced with cholesterol in disordered phases [32–35]). That is, the timescale is unlikely to be due to a contaminant or oxidated lipid. The timescale appears too small to be a result of lipid flipflop, whose characteristic timescale is on the order of hours [36]. Although the mechanism for the small amplitude slow relaxation in pure POPC is unknown, the effect is larger in PE/PC. While Δ*c*_PE_ for PE/PC and POPC (0.340 nm^−1^ and 0.16 nm^−1^) appear comparable, the strength of the effect goes as 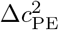, and thus is over four times larger for PE/PC.

### D. Implication of diffusional softening

Fitting the distribution of undulation *magnitudes* is sensitive to *κ*_apparent_. Fitting undulation *kinetics* extracts *κ*, *D* and 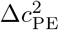. As presented here, *κ* only accounts for diffusional softening; structural dynamics that couple to curvature at faster timescales affect *κ*.

Kinetic fitting of GUVs demonstrates that the PE/PC mixture has *κ*, *D* and Δ*c*_PE_ consistent with previous measurements of spontaneous curvature [30] and diffusion [33], as well as a minimal difference if any, for *κ* relative to POPC. That is, the larger and somewhat slower undulations of the PE/PC mixture are consistent not with a change in the underlying softness of the bilayer, but rather through the coupling of the spontaneous curvature of DOPE to dynamic undulations.

Separating mechanistic contributions to *κ*_apparent_ is critical for developing a complete model of the membrane, including its equilibrium and non-equilibrium relaxation behavior. In the absence of the model of diffusional softening, extrapolating *κ*_apparent_ to 100% DOPE yields, relative to pure POPC, an extremely small value for pure DOPE, ca. 10.64 *k*_B_*T*. In contrast, in the diffusional softening model, *κ* depends weakly on DOPE fraction. Note that while pure DOPE does not readily form lamellar phases at this temperature, its bending modulus in the hexagonal phase is 22-26*k*_B_*T*, depending on the incorporation of interstitial tetradecane [30].

The principal indication of diffusional softening is the long-timescale auto-correlation of the undulation amplitude. The initial fast undulation kinetics cannot be used reliably to determine *κ*; the undulation rate *k*_+_ is reduced by coupling to diffusion (by 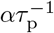). The impact of this “friction” is reduced at high *q* as *τ*_p_ grows relative to *τ*_m_.

### E. GUV relaxation kinetics as a probe for structural heterogeneity

Any structure with curvature coupling different from the bulk will influence the magnitude and kinetics of GUV fluctuations. Thus, in theory, kinetics can be used to infer the dynamics and coupling strength of complex structures such as nanodomains and lipid multimers. Coupling of GM1 to curvature is a plausible explanation for the dramatic softening of POPC/GM1 mixtures [37], where at mol fractions less than 10% GM1 the bending modulus is less than 25% of that of pure POPC. Such softening could indicate that size (*A*_p_) and/or coupling strength (Δ*c*_PE_) of the GM1-enriched structural unit exceeds that of a typical lipid. The proximity of the gel/fluid transition suggests the possibility of curvature-sensitive GM1 multimers.

Experiments clearly indicate liquid ordered domains have increased *κ* compared with disordered phases [7]. The magnitude of the effect is likely sufficiently large that the linearized treatment of the coupling *α* is inadequate. For stiff domains that couple strongly to curvature, a quickly becomes larger than one, indicating a breakdown in the theory. Indeed softening is non-linear above 5 mol% GM1 in POPC.

## IV. CONCLUSION

The dynamic coupling of the lateral distribution of curvature sensitive lipids to membrane undulations leads to diffusional softening of the membrane [12, 15, 18, 21]. The undulation autocorrelation function implies the intrinsic bending rigidity of the membrane, the diffusion constant of the underlying lipids, and the magnitude of the spontaneous curvature difference (through the timescale). The intrinsic bending rigidity of the membrane determined from the kinetics is related to the apparent bending rigidity determined from the fluctuation spectrum through the softening factor. The experiment and model corroborate a similar experiment on membrane nanotubes by Bashkirov et al [21], in which DOPE also softened a majority PC bilayer according to Eq. 25.

A key factor of the *diffusional softening* mechanism is how coupling of undulations to diffusion acts as a “friction”. The impact of this force depends on *q*. The response of membrane undulations to this frictional force is observed in the relaxation kinetics of membrane undulations. Membrane viscosity [38] and interleaflet friction [39, 40] also influence membrane undulation kinetics at shorter timescales. Understanding how undulation kinetics is modified by these terms has proved crucial for understanding the fine mechanisms of membrane shape dynamics [41].

It is critical to understand the extent of this mechanism when inferring the bending modulus of complex mixtures. For example, conflicting results have recently been published for the bending modulus of cholesterol in DOPC. Neutron spin-echo experiments, which probe relaxation times of bilayers below the timescale of diffusion, indicate the bilayer is stiffer [42]. Yet multiple techniques that probe equilibrium fluctuations have shown that *κ*_apparent_ is unchanged [8, 19, 29, 43]. Hexagonal phase experiments indicate that cholesterol will have a high negative spontaneous curvature in fluid bilayers [30]. Accepting the diffusional softening mechanism, cholesterol is expected to *soften* a DOPC bilayer, in the absence of a stiffening effect. Note however, that to fully resolve the reported discrepancy, membrane viscosity contributions in time-correlation analysis should be interrogated; recent analysis indicates that viscosity affects a broader array of undulations of smaller liposomes than previously anticipated, suggesting another possible change in the interpretation of spin echo experiments [44].

GUV fluctuations suggest that cholesterol at 10 mol% reduces the *κ*_apparent_ of SOPC, while increasing *κ*_apparent_ at higher concentrations [45]. X-ray derived data contradicts this [43]. In either case the effect is sufficiently small as to be difficult to statistically distinguish. Hexagonal phase experiments with variable osmotic stress (a technique for which lateral redistribution is irrelevant) indicate cholesterol stiffens DOPC and DOPE somewhat. Considering diffusional softening, it is possible that cholesterol is both stiffening the underlying *κ* while *lowering *κ**_apparent_ such that the change is minimal. This case illustrates the importance of deducing the molecular mechanism of changes in bilayer stiffness.

## V. ACKNOWLEDGMENTS

K.S. and A.J.S. were supported by the intramural research program of the Eunice Kennedy Shriver National Institutes of Child Health and Human Development (NICHD) at the National Institutes of Health. M.A. acknowledges funding from International Max Planck Research School on Multiscale Bio-Systems (IMPRS). This research was also supported in part by the National Science Foundation under grant NSF PHY-1748958.

## Notes

https://figshare.com/articles/media/GUV_mp4/21224636

### Competing Interest Statement

The authors have declared no competing interest.

