## Supplemental Material for "Kinetic relaxation of giant vesicles validates diffusional softening in a binary lipid mixture"

Kayla Sapp,[1] Mina Aleksanyan,[2,3] Kaitlyn Kerr,[1] Rumiana Dimova,[2] and Alexander Sodt[1]

[1]*Eunice Kennedy Shriver National Institute of Child Health and Human Development, National Institutes of Health, Bethesda, Maryland*

[2]*Max Planck Institute of Colloids and Interfaces, 14476 Potsdam, Germany*

[3]*Institute for Chemistry and Biochemistry, Freie Universität Berlin, 14195 Berlin, Germany*

#### A. Experimental Fluctuations

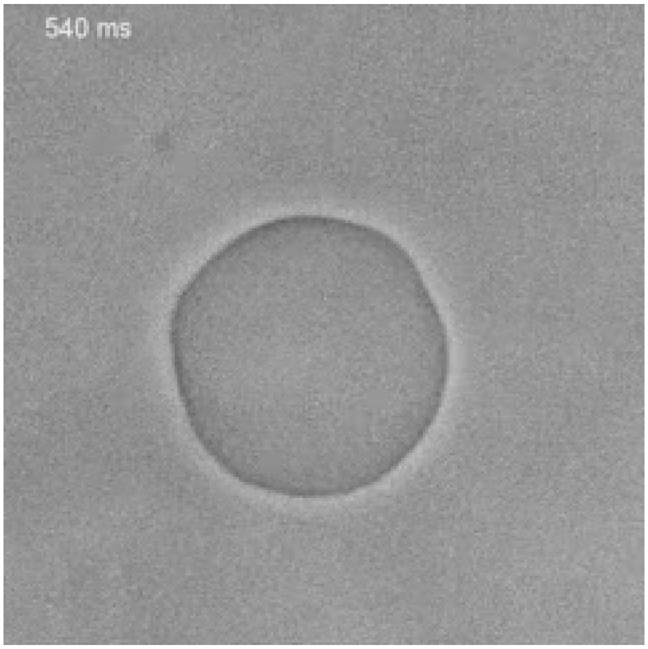


FIG. S1: A slowed-down video of a fluctuating POPC vesicle containing 40 mol% DOPE (same vesicle as in Figure 1 in the main text). The vesicle was prepared in 20 mM sucrose, 4-fold diluted in 22 mM glucose, and additionally deflated by leaving the observation chamber open for 5 minutes. The sequence shows phase contrast images acquired at 100 frames per second and displayed at 25 frames per second (processing was done with Fiji). The approximate duration is roughly 4 seconds in real time (time stamps shown in the upper left corner). (DOI: 10.6084/m9.figshare.21224636)

#### B. Sensitivity to fit choices

Figure S2 shows the modest variation in apparent spontaneous curvature difference with $q_{\max}$.

Figure S3 shows the variation in the apparent spontaneous curvature difference with $t_{\max}$. The fit parameters

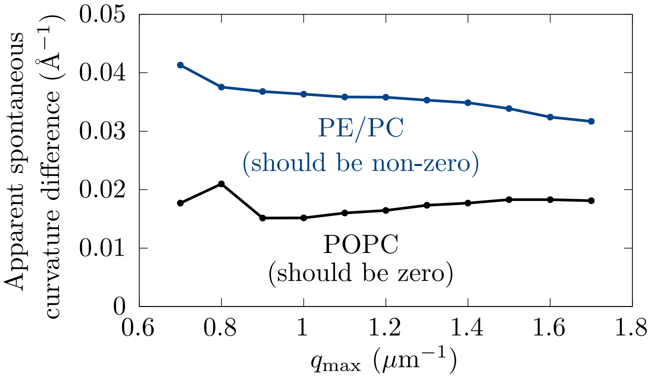


FIG. S2: Variation in the apparent spontaneous curvature difference with $q_{\max}$.

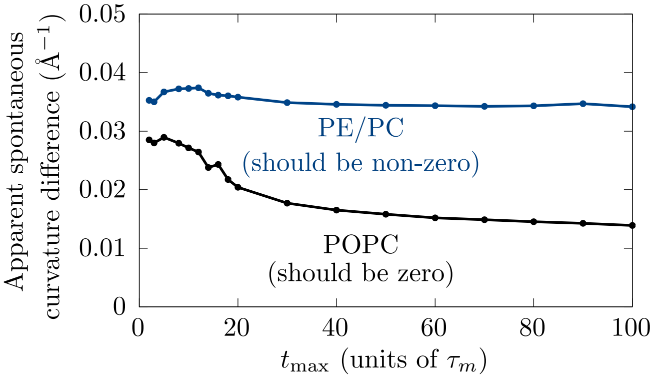


FIG. S3: Variation in the apparent spontaneous curvature difference with $t_{\max}$.

converge only when the time-domain is extended well beyond that of the membrane relaxation.

### I. COMPLETE SET OF FIT HISTOGRAMS

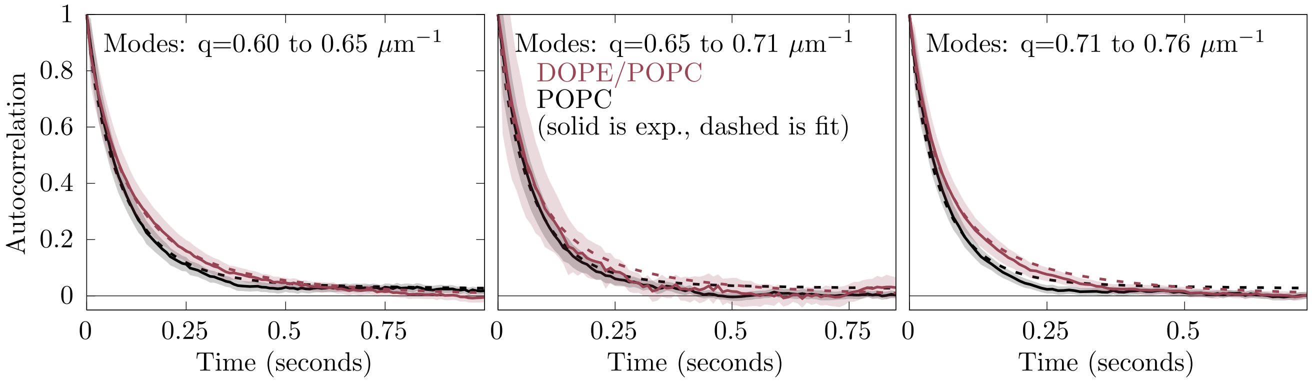


FIG. S4: The average of the autocorrelation $\langle \nu_q(t)\nu_q(0)\rangle$ over similar modes for the experiment (solid) and fits (dashed). POPC is colored in black. PE/PC is colored red. Filled curves indicate two standard errors obtained by averaging over similar modes.

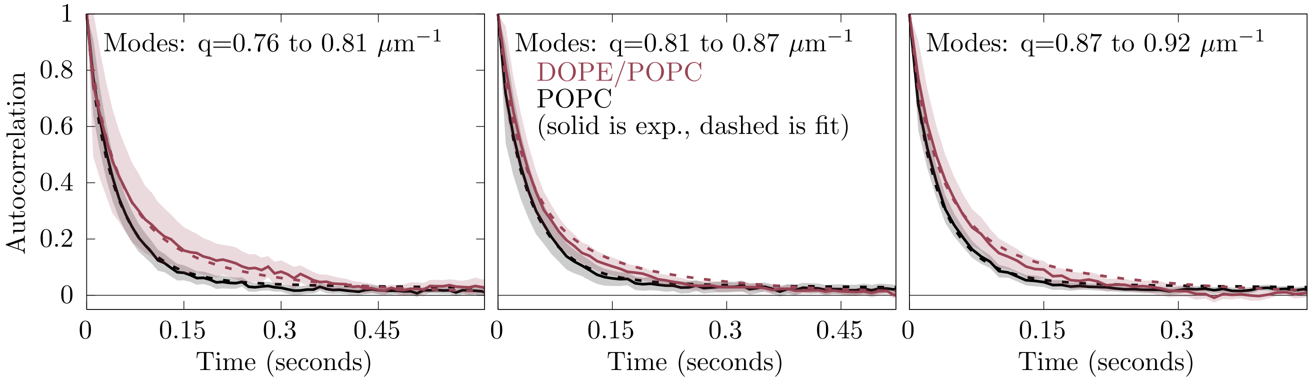


FIG. S5: The average of the autocorrelation $\langle \nu_q(t)\nu_q(0)\rangle$ over similar modes for the experiment (solid) and fits (dashed). POPC is colored in black. PE/PC is colored red. Filled curves indicate two standard errors obtained by averaging over similar modes.

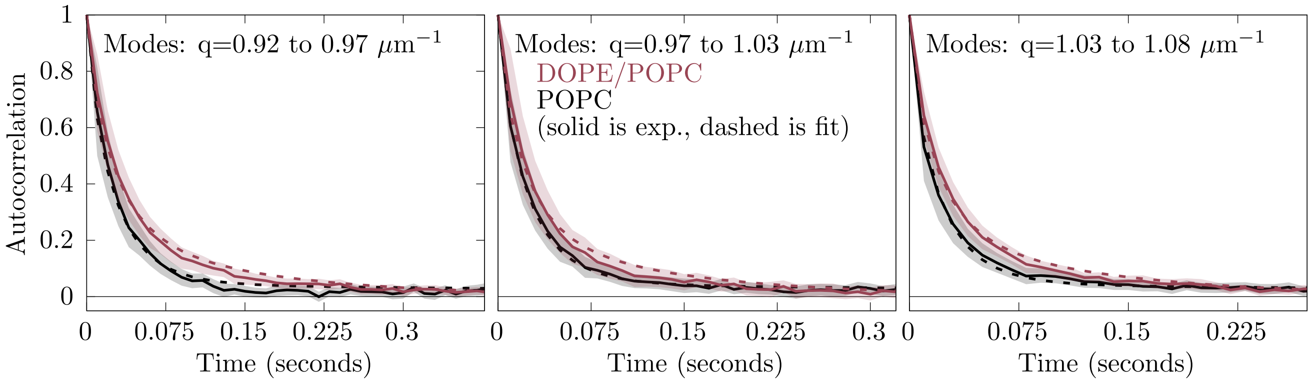


FIG. S6: The average of the autocorrelation $\langle \nu_q(t)\nu_q(0)\rangle$ over similar modes for the experiment (solid) and fits (dashed). POPC is colored in black. PE/PC is colored red. Filled curves indicate two standard errors obtained by averaging over similar modes.

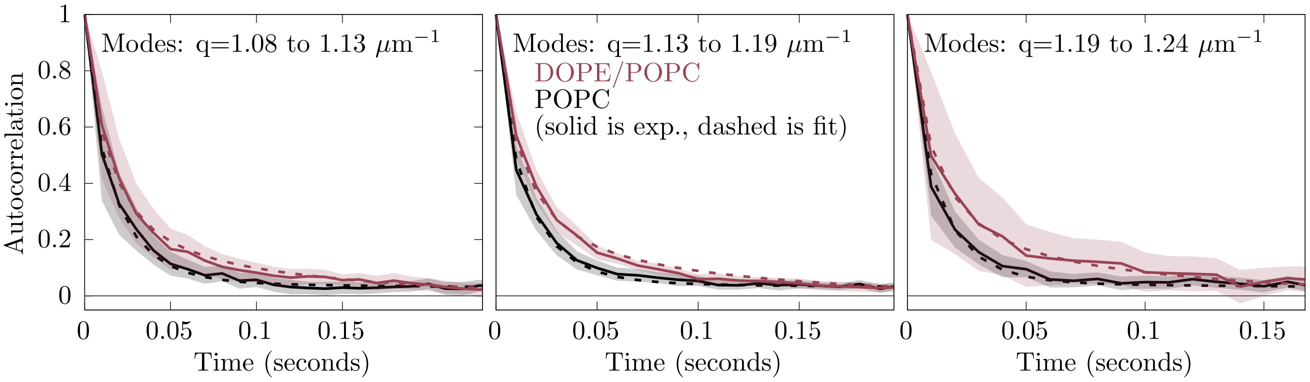


FIG. S7: The average of the autocorrelation $\langle \nu_q(t)\nu_q(0)\rangle$ over similar modes for the experiment (solid) and fits (dashed). POPC is colored in black. PE/PC is colored red. Filled curves indicate two standard errors obtained by averaging over similar modes.

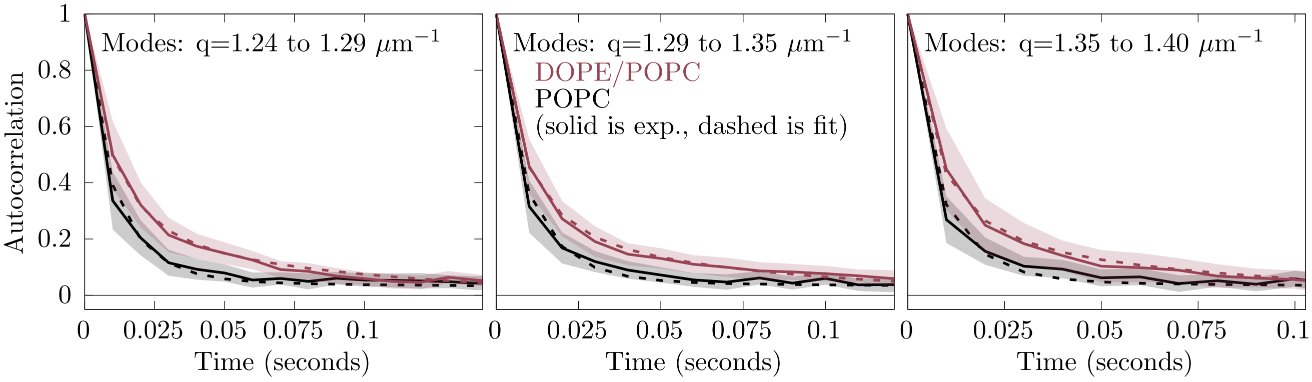


FIG. S8: The average of the autocorrelation $\langle \nu_q(t)\nu_q(0)\rangle$ over similar modes for the experiment (solid) and fits (dashed). POPC is colored in black. PE/PC is colored red. Filled curves indicate two standard errors obtained by averaging over similar modes.
